# Characterizing the Non-Normal Distribution of Flow Cytometry Measurements from Transiently Expressed Constructs in Mammalian Cells

**DOI:** 10.1101/057950

**Authors:** Peter F. McLean, Christina D. Smolke, Marc Salit

## Abstract

In mammalian cells, transient gene expression (TGE) is a rapid, minimal-investment alternative to single-copy integrations for testing of transgenic constructs. However, transient gene expression, as measured by flow cytometry with a fluorescent reporter, typically displays a broad, asymmetric distribution with a left-tail that is convolved with background signal. Common approaches for deriving a summary statistic for transiently expressed gene products impose a normal distribution on gated or ungated data. Summary statistics derived from these models are heavily biased by experimental conditions and instrument settings that are difficult to replicate and insufficient to accurately describe the underlying data. Here, we present a convolved gamma distribution as a superior model for TGE datasets. The 4-6 parameters of this model are sufficient to accurately describe the entire, ungated distribution of transiently transfected HEK cells expressing monomeric fluorescent proteins, that operates consistently across a range of transfection conditions and instrument settings. Based on these observations, a convolved gamma model of TGE distributions has the potential to significantly improve the accuracy and reproducibility of genetic device characterization in mammalian cells.

## Introduction

Notwithstanding an incomplete understanding of mammalian *in vivo* gene expression, many expression products measured by flow cytometry can be reliably represented by well-described distributions. For example, stable integrations of transgenic fluorescent reporters under the control of constitutive promoters often give rise to an approximately lognormal fluorescence signal distribution. A faster and less expensive alternative to integrated, single-copy gene expression is transient gene expression (TGE), making it an attractive method for rapid prototyping of genetic constructs with minimal investment.

However, transient gene delivery introduces a wide range of plasmid copy numbers that will degrade and dilute over time^1–3^. The resulting distributions of measured fluorescence intensities from transiently expressed constructs are generally non-normal, span multiple decades, and have non-negligible overlap with background signal. In light of these asymmetries and uncharacterized distribution parameters, careful consideration should be given to the model used to generate a summary expression statistic from the measured signal distribution. Notwithstanding, typical analyses of TGE experiments assume one or two normal distributions on gated or ungated fluorescence measurements, respectively, which fail to accurately represent the data (Fig. 1). Critically, gaussian-based models only attempt to estimate a mean signal (lacking reproducible information on other characteristics of the underlying distribution) and are heavily influenced by background signal. Given that the background signal and the ratio of signal-to-background are sensitive to experimental conditions, technical configurations, and arbitrary instrument settings, the results of TGE experiments are often difficult to replicate.

**Figure 1:**
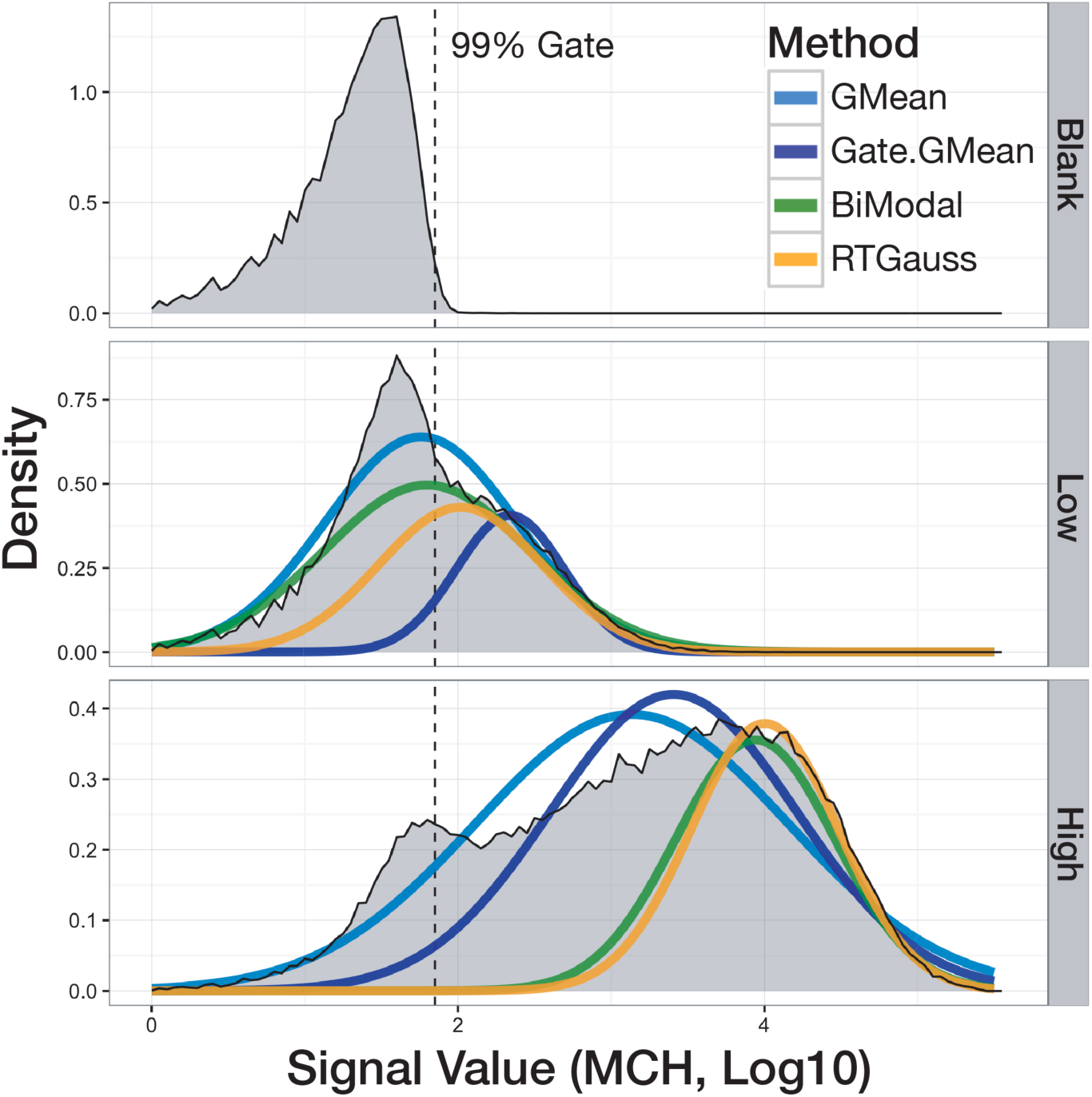
Common Gaussian-based Models for TGE Analysis. The distribution of HEK cells transiently expressing mCherry is broad and asymmetric; common gaussian-based models are poor descriptors of the actual distribution. Data shown: null transfection (Blank), qualitative low and high transfected DNA concentrations (Low, High). Methods: geometric mean of the whole dataset (GMean), geometric mean of the gated dataset (dashed line, Gate.GMean), the upper component of a bi-modal distribution fit to the log-transformed dataset (Bimod), a gaussian curve fit to the right-tail of the log-transformed dataset (RTGauss).

In contrast, the convolved gamma model here presented parameterizes the entire, ungated distribution and generates 4-6 parameters that are sufficient to accurately reconstruct the entire dataset, i.e., the dominating components of noise, background, and target fluorescence. We also demonstrate how the gamma parameters provide a robust measure of mean gene expression across at least 3.5 orders-of-magnitude and in samples with very low signal-to-background ratios. We further posit that the gamma parameters encode information on transfection efficiency that is not biased by arbitrary instrument settings and have potential applications for characterizing the transcriptional and translational states of the cells.

## Methods

### Cell Transfections

Two fluorescent reporters, tagBFP (BFP) and mCherry-PEST (MCH; C-terminal PEST degradation sequence), were co-expressed from separate plasmids by CMV promoters in HEK 293 FlipIn cells (ThermoFisher). For the concentration series, the Mirus transfection reagent (REF) was used to co-transfect BFP at a constant concentration of 15 fmol per well and MCH was introduced in half-log increments from .015 fmol to 150 fmol per well. For the transfection reagent series, we used Lipfectamine 2000 (REF), Fugene HD (REF), Mirus (REF), and TurboFect (REF). Empty vector was used to raise all sample mixes to a constant DNA mass. In most cases, transfection mixes were constructed such that each well was transfected with 600 ng of total DNA and 1.8 μL of transfection reagent (3 μL:1 μg), as is manufacturer recommended. TurboFect was mixed with 1.2 μL reagent (2 μL:1 μg). All co-transfections were conducted in duplicate in 24-well plates into which 150,000 cells were seeded 24 hours prior. Samples were incubated for 48 hours with the applied transfection mix and measured on a Miltenyi Biotec MACSQuant VYB cytometer with 405 nm and 561 nm excitation, and 450 nm/±50 nm and 615 nm/±20 nm filters, for BFP and MCH, respectively.

### Data Processing

All flow cytometry data processing was conducted in R, with the aid of the following packages: flowCore, flowViz, flowBeads, ggplot2, reshape2, Hmisc, RColorBrewer, modeest, stats4, plyr, msm, distr, MASS, and mixtools (flow* packages courtesy of BioConductor.org). Briefly, data was imported as FCS files and events were filtered by way of a density-based forward/side scatter gate and a forward-scatter area/height gate; when combined, gates retained ∼70% of measured events (representing 56,000–62,000 events per sample). Each replicate contained null-transfected and single-color control samples that were used to calculate a compensation matrix for spectral correction. Ultimately, no spectral compensation was used for the BFP/MCH fluorophore pair (overlap < 0.5%). All saturated events were dropped. Raw MCH values were converted to “equivalent BFP” values (denoted “eqMCH”) using a linear-regression model fit to the log-transformed values of cells expressing equal amounts of both plasmids—a necessary step in order to justify direct inter-channel comparisons and arithmetic. An additional benefit of the described inter-channel conversion is that samples acquired under different detector amplification settings (gains) are converted to the same scale, i.e., equivalent BFP signal value. Parameterization was conducted separately for all replicates.

## Results

### Characterization of Background Signal

The necessary first step in developing an ungated model for TGE flow cytometry data is to define a model for the background signal. This was accomplished by measuring null-transfected cells with increasing gain values and testing the fit of various known distributions to the sample data (Fig. 2). At low gain, events are normally distributed around a value close to zero; the negative values arise from the combined effects of imperfect baseline-restoration in the detectors and various other contributions of linear measurement noise. With increasing amplification, the background signal is dominated by multiplicative cellular autofluorescence and is modeled as a lognormal distribution. The transition between the two distributions is accurately modeled as a convolution of the two functions, namely, normal and lognormal distributions. Thus, the description of background signal with either two parameters (normal or lognormal) or four parameters (convolved normal and lognormal) needs to be determined for each experiment and instrument configuration.

**Figure 2:**
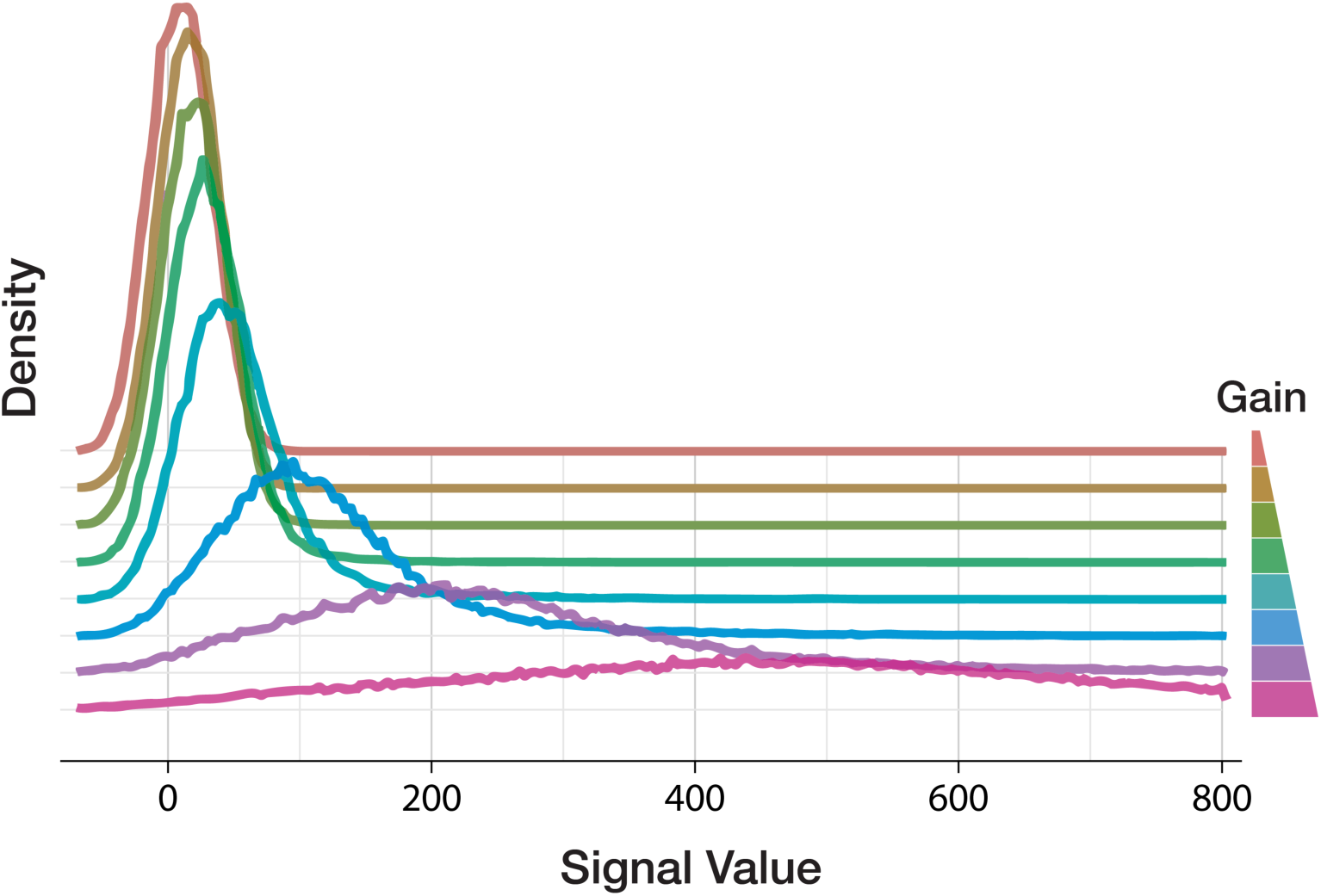
Determination of Background Signal as a Two-Component Convolution. Measurement of null-transfected HEK cells with increasing the detector amplification values (gain) demonstrates the transition from a linear, normally distributed background at low gain values to an exponentially distributed, lognormal distribution at high gain values. The data is well-described by a convolution of the two functions across the transition. Gain values are represented qualitatively because they are instrument specific.

(Note: instruments with log-amplifiers, as opposed to the linear-amplifiers in the instrument that was used for these experiments, generate data that is truncated at zero; low-level background signal could be modeled with truncated distributions as necessary.)

### Convolved Gamma Distribution

Of the exponential family distributions we tested, the gamma distribution best resembled the asymmetries observed in TGE data. Importantly, the use of a gamma distribution, Gamma(*kθ*), to describe gene expression is not without precedent^4^. The complete model is therefore composed of a convolved normal, lognormal, and gamma distribution, and represents all sources of observed single in TGE flow cytometry data, i.e., the linear noise/background component, the lognormal background component (autofluorescence), and the gamma-distributed fluorescent reporter signal.

Owing to the large range of signal values, sparse distribution of high-value events, and existence of negative-value events, numerical-integration and maximum-likelihood estimation (MLE) approaches for generating the full convolved distribution and parameterizing datasets were frequently unstable, non-convergent, and computationally impractical. However, convolving only the background components (normal and lognormal) within typical experimental ranges is relatively simple using a fast-Fourier transform algorithm from the “distr” R package; this offers a fast method for parameterizing the background signal of an experiment. Unfortunately, this approach is not successful when including the gamma component. Instead, a simulation-based approach is used in which the contributing distributions are sampled and added to generate a simulated convolved dataset from a set of start values. The simulated dataset is then log-transformed, binned, and used to fit parameters for the similarly log-transformed and binned sample data using a least-squares method. To simplify parameterization of the gamma component, the four parameters of the convolved background signal can be derived from null-transfected or non-expressing control samples and fixed for subsequent parameterizations for all samples with similar background characteristics (e.g., unique cell-type, single batch) with little impact on the resulting gamma parameter values.

The resulting parameter values can then be used to accurately simulate the entire, ungated TGE distribution (Fig. 3) for a wide range of transfected DNA concentrations and across multiple instrument configurations. As discussed previously, in circumstances of very low autofluorescence or high amplification, it may also be appropriate to model background with either the normal or lognormal parameters, resulting in final model consisting of only four parameters—two of which (describing background) can be independently measured in control samples.

**Figure 3:**
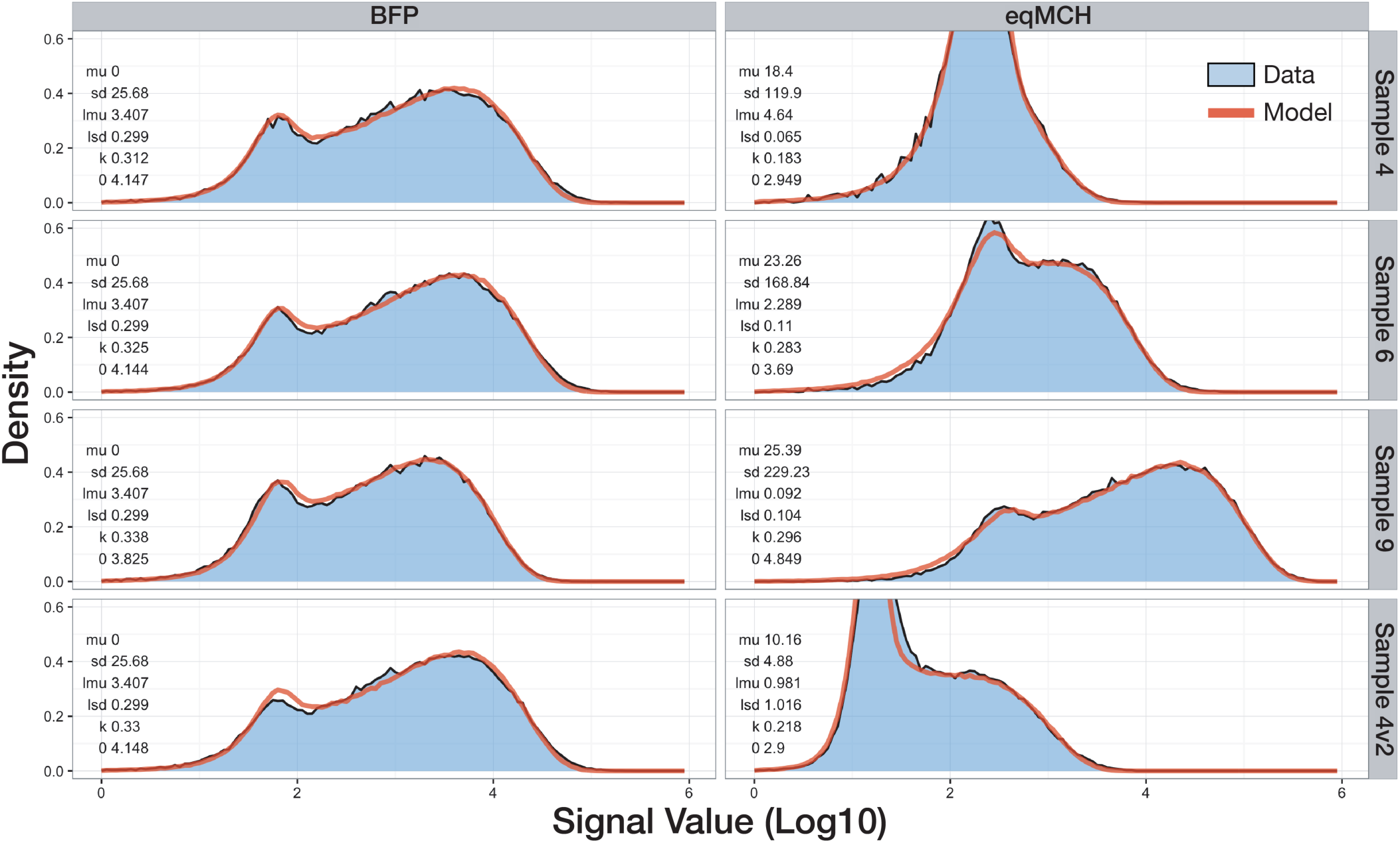
Convolved Gamma Model Describes the Full TGE Distribution. Representative samples from a transfection concentration series reflecting low, medium and high expressing populations are very well described by both a two-parameter fit (fixed background parameter values, BFP) and a six-parameter fit (eqMCH). Additionally, samples measured at different PMT voltages (gains) are equivalently parameterized by the model (Sample_04v2). Sample data represents the sum of two replicates. Parameter values are given in each panel: N(mu, sd), logN(lmu, lsd), G(k, theta).

Together, the shape and scale parameters of the gamma distribution are sufficient to reconstruct the fluorescence distribution absent the contribution of background signal. As such, the gamma-derived mean serves as a summary statistic that faithfully represents the underlying data. At a minimum, the gamma parameters allow a systematic determination of transfection efficiency that is not biased by background signal, and reveal a concentration-dependent limit to the shape parameter that exists two orders-of-magnitude below manufacturer-recommended transfection concentrations (Fig. 4). Importantly, samples are equivalently parameterized at different detector amplification settings, supporting the use of a convolved gamma distribution for reproducibly modeling TGE data collected with arbitrary instrument settings.

**Figure 4:**
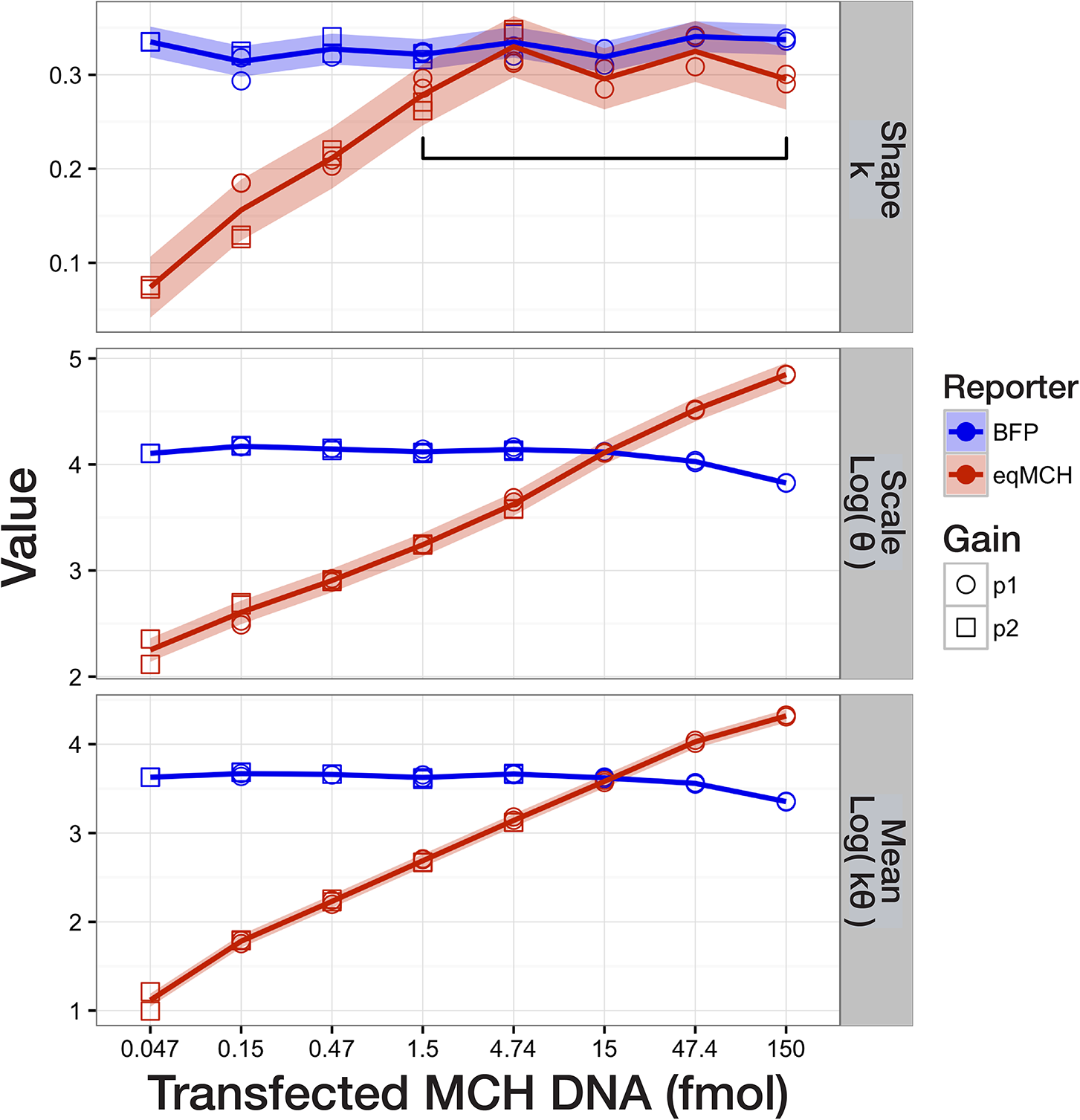
Analysis of Gamma Parameters. Fitted parameters for a transfection concentration series measured at two gain values (MCH only) reveal important characteristics of the distributions. Shape, scale, and derived mean values are equivalently parameterized between samples measured at different gain values. Scale and mean parameters are log-transformed for visualization only; mean is calculated as E[*X*]=*kθ*. Shaded ribbon indicates ±2 standard deviations of the pooled variance (log) for each parameter and reporter.

### Method Comparison

We can evaluate the performance of the various models by investigating the relationship between the amount of transfected DNA and the estimated mean fluorescence signal. In contrast to bacterial systems where ribosome availability appears to be the primary bottleneck for gene expression, mammalian transgenic gene expression is believed to operate far from the metabolic limit of the cells^5–7^. Under this hypothesis, we should expect a near-linear relationship between the amount of transfected DNA and the measured gene expression; deviating only at the high transfection concentrations and in cells with high gene expression (Fig. 5A). Furthermore, co-transfected plasmids are presumed to operate non-interactively, competing equally for resources only in circumstances of very high expression. This would manifest as slight negative-curvature in a plot of mean fluorescence and summed DNA (Fig. 5B), and a 1:1 relationship between the ratios of transfected plasmids and of fluorescence means of two reporters (Fig. 5C). Thus, the various analysis models can be evaluated by their ability to generate reproducible mean estimates across a range of transfection conditions and instrument settings and in their concordance (expected linearity) with current knowledge of mammalian transgenic gene expression.

**Figure 5:**
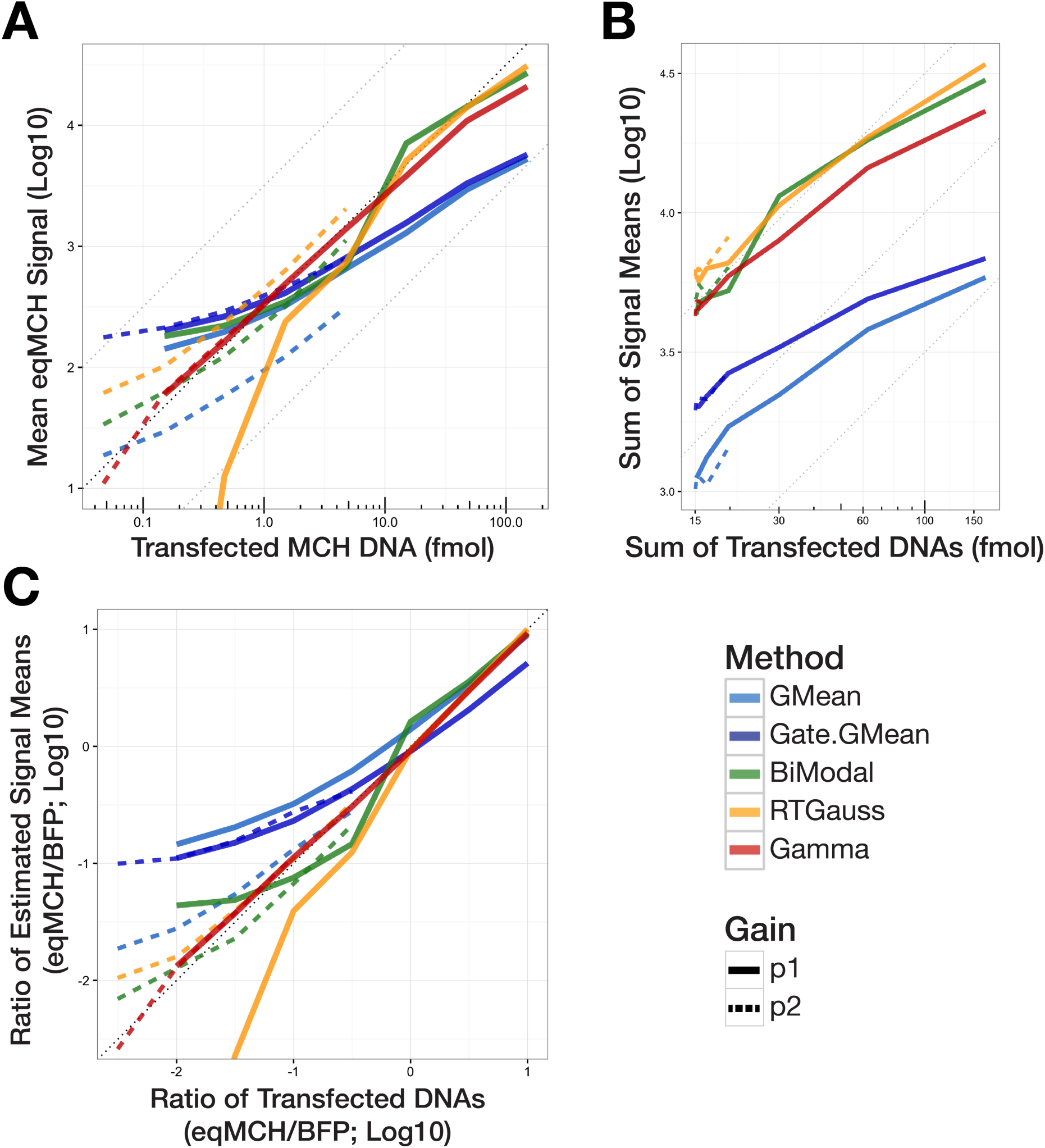
Method Comparison. The estimated means of five analysis models were determined for a transient transfection concentration series in HEK cells and plotted against the amount of transfected DNA. MCH was measured with two gain values (solid or dashed). Estimated means represent the average of two replicate experiments. A) MCH alone; B) the sum of MCH and the BFP control; C) the ratio of MCH and the BFP control.

The simplest approach for deriving a summary statistic is to calculate the geometric mean of the gated or ungated data. A more advanced model is that of a bi-modal distribution, which is typically applied to ungated data and makes the assumption that both the background and target fluorescence signal distributions are normally distributed. Fitting a gaussian curve to the right-tail of the TGE distribution (RTGauss) provides a measure of target fluorescence similar to the bi-modal distribution, but which excludes events at the level of background signal and is more consistent in samples with low signal-to-background ratios as a result.

The described gaussian-based and the convolved gamma model proposed here were used to analyze a transfection concentration series in which MCH was transfected over a wide range of concentrations and BFP was held constant. For all samples, MCH fluorescence was additionally measured with two detector gain values (Fig. 5). As anticipated, the means of the uni-modal gaussian models, whether gated or ungated, are strongly influenced by their position relative to background. This is observed both as a horizontal asymptote at the level of background signal and as a mean signal shift between the samples measured at different gain values (Fig. 5A-C). This occurs because using ungated data gives equal weight to background signal and target fluorescence, while gated data typically results in a severely truncated distribution.

The bi-modal model provides a more consistent description of the mean target signal by circumventing some of the background signal biases by using ungated data. It is nonetheless unpredictably sensitive to changes in the shape of the underlying distribution that can result from changing instrument settings and transfection reagents, and is ultimately a poor descriptor of both the background signal and target fluorescence distributions (Fig. 1). The RTGauss-derived mean is more robust to low signal-to-background ratios, but also appears to be biased in these samples (diverging results at different gains, Fig. 5A,C). Of these approaches, only the bi-modal model explicitly accommodates the background signal with which target fluorescence is convolved, but offers no useful characterization of the background signal distribution—using it only as a placeholder for determining the position of the upper (target signal) distribution.

In contrast, the gamma-derived mean operates consistently across gain settings and over a range of at least 3.5 orders-of-magnitude. As expected, we observe a near-linear relationship between transfected MCH plasmid and mean MCH fluorescence signal and an equal, linear relationship between the ratios of co-transfected plasmids and fluorescence means (Fig. 5A,C).

### Impact of Transfection Reagents

It is typically difficult to compare TGE experimental results conducted with different transfection reagents because they result in distributions with slightly different characteristics; existing, gaussian-based models incompletely characterize these differences (Fig. 1). In contrast, the convolved gamma model accurately describes several different lipid-based transfection reagents—making possible an evaluation of distribution characteristics (Fig. 6). Importantly, this allows for a systematic determination of transfection efficiency that is based on the structure of the distribution rather than on the percent of cells measured above background (which is subject to the effects of arbitrary instrument configuration).

**Figure 6:**
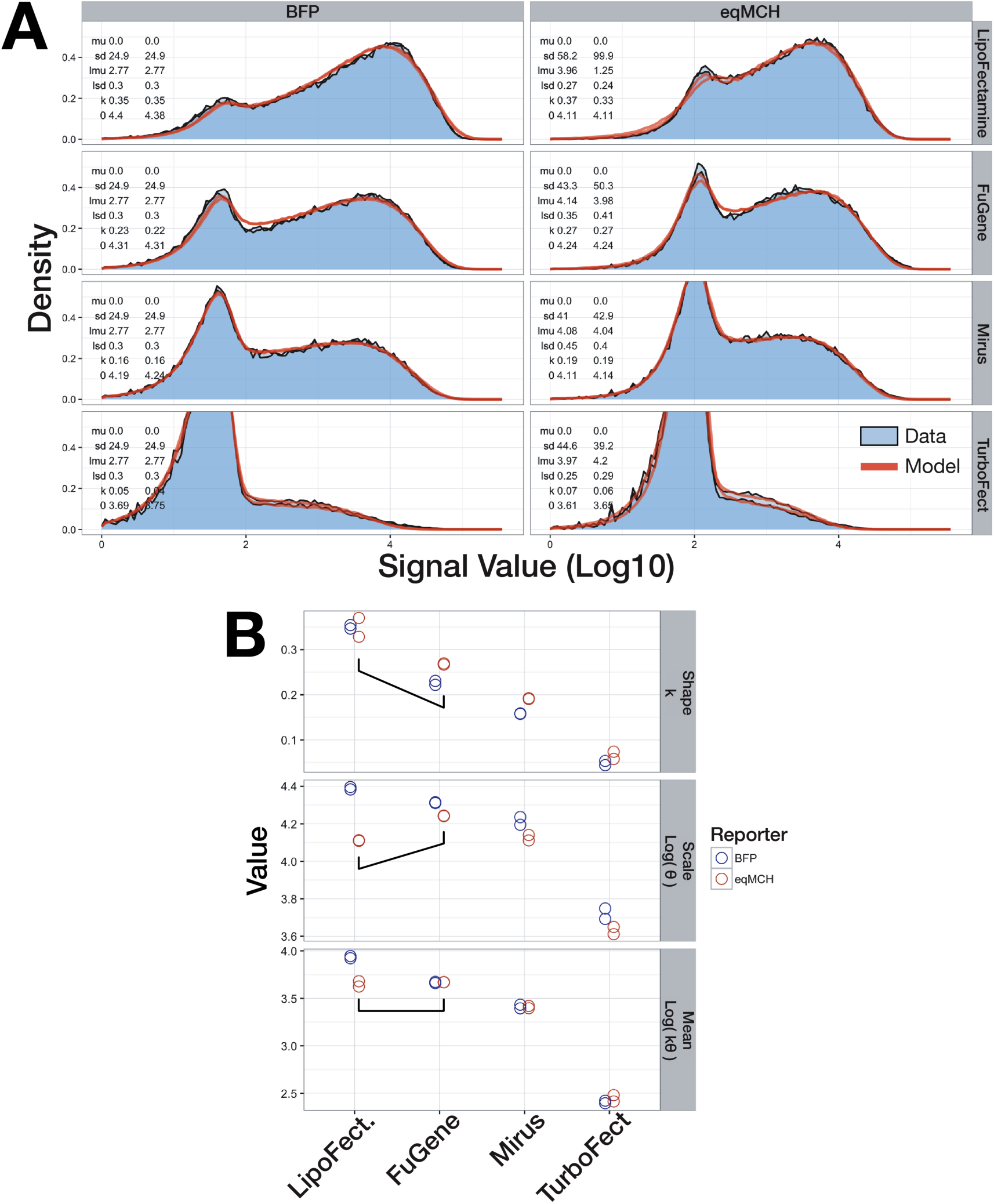
Evaluating the Effects of Transfection Reagent on TGE Distributions with the Convolved Gamma Model. A) The convolved gamma model was found to accurately describe HEK cells transfected with four different lipid-based transfection reagents. Shown are the data and fitted curves for two replicates. Parameter values are given in each panel: N(mu, sd), logN(lmu, lsd), G(k, theta). NOTE: Transfection conditions were not optimized for each reagent, and should not be interpreted as indicative of reagent quality. B) Evaluation of the fitted gamma parameters shows some degree of differential effects on shape and scale between reagents (highlighted).

## Discussion

Transient gene expression (TGE) is an important, and widely used method for rapidly testing genetic constructs in mammalian cells^8^. Naïve analysis of TGE flow cytometry data commonly assumes normally distributed data and estimates only a position parameter (mean), offering no meaningful measure of any other distribution characteristic. In contrast, a convolved gamma distribution accurately describes the entire, ungated distribution of transiently expressed genes in mammalian cells with 4-6 parameters. As such, the convolved gamma distribution helps to resolve existing ambiguities in mammalian TGE behaviors and was shown to be more robust across a larger range (at least 3.5 orders-of-magnitude) of experimental conditions and instrument settings than other methods of analysis in this study.

Importantly, the gamma parameters offer information on TGE distributions that are not accessible with gaussian-based models such as transfection reagent-specific characteristics. The gamma parameters may also reflect underlying biophysical processes. For example, Freidman *et.al*.^4^ demonstrates a link between the shape and scale parameters of the gamma distribution to the cellular states of transcription and translation, respectively, in bacterial cells; the shape describing the frequency of protein production events and the scale reflecting the amount of protein produced per event. However, given important differences in the maturation of transcripts, dynamics of transcription and translation, and gene copy number between bacterial and mammalian systems, a biophysical rationale for the application of a gamma model to mammalian gene expression has yet to be developed.

Nonetheless, the gamma parameters have clear empirical value in providing an accurate description of the fluorescence signal distribution and in characterizing observed experimental effects. For example, the observed plateau that occurs for the shape parameter in the transfection concentration series (Fig. 4) may be indicative of a saturation effect on one or more of the mechanisms that mediate DNA delivery, e.g., escape from the endosomes, cytoplasmic concentration during mitosis, or nuclear shuttling activities^5,7,9–13^. Alternatively, it might reflect a transition in the dominating source of noise, where stochastic expression effects begins to dominate over gene-delivery variance. Future experiments might derive additional utility of this model by identifying the contribution of various transfection reagents (i.e., the distribution of transcription-competent DNA in the nuclei) to the observed gamma distribution, enabling a reagent-specific deconvolution of copy-number from the full distribution, and ultimately, a characterization of transgenic gene expression independent of the mechanism of its delivery.

Thus, the convolved gamma distribution offers better representation of TGE that can improve the accuracy and reproducibility of genetic device characterization in mammalian cells. More broadly, more accurate methods of analysis can help standardize the characterization of biological parts, ultimately supporting the establishment of a viable synthetic biology ecosystem for mammalian systems.

## Acknowledgements

We also thank NIST colleagues Ariel Hecht, Matt Munson, Sarah Munro, and David Ross, as well as those of the Smolke lab at Stanford University for their advice and support.

## Disclaimer

Certain commercial equipment, instruments, or materials are identified in this paper in order to specify the experimental procedure adequately. Such identification is not intended to imply recommendation or endorsement by the National Institute of Standards and Technology (NIST), nor is it intended to imply that the materials or equipment identified are necessarily the best available for the purpose.

